# A workflow for incorporating cross-sectional data into the calibration of dynamic models

**DOI:** 10.1101/2023.01.17.523407

**Authors:** Sophie Fischer-Holzhausen, Susanna Röblitz

## Abstract

Mathematical modelling and dynamic simulations are commonly used in systems medicine to investigate the interactions between various biological entities in time. The level of model complexity is mainly restricted by the number of model parameters that can be estimated from available experimental data and prior knowledge. The calibration of dynamic models usually requires longitudinal data from multiple individuals, which is challenging to obtain and, consequently, not always available. On the contrary, the collection of cross-sectional data is often more feasible. Here, we demonstrate how the parameters of individual dynamic models can be estimated from such cross-sectional data using a Bayesian updating method. We illustrate this approach on a model for puberty in girls with cross-sectional hormone measurement data.

## 1 Introduction

Mechanistic models formulated as systems of ordinary differential equations (ODE) are widely used to study the dynamic behaviour of biological systems (Motta and Pappalardo, 2013). Often, only a limited number of model parameters are known, and at least a subset of the model parameters must be estimated to construct a useful model. The model calibration (parameter estimation) is typically done by fitting the model to experimental data, which poses several challenges (Gábor and Banga, 2015). In systems biology, we often encounter parameter landscapes with multiple local optima, complicating the parameter optimization. Additionally, there are different sources of uncertainty, such as a lack of prior knowledge, noisy data, ambiguities in the model structure and a mismatch between model complexity and available data. Despite those challenges, model calibration and analysis are central to creating a meaningful model (Villaverde et al., 2022). For the calibration of a dynamic model, one needs information about the system’s time evolution (longitudinal data/time-series data). In general, data collection is time and resource-consuming, and the collection of time-series data comes with additional challenges (Udtha et al., 2015). Especially in clinical studies, the acquisition of longitudinal data requires a long-term commitment of study participants (Robinson et al., 2007), and the number of samples that can be taken is limited. Consequently, the data sets are sparse in the number of time points, and the sample size tends to be small (Greenland et al., 2016). In addition, populations often show a high inter- and inter-individual variability (Sebastian-Gambaro et al., 1997; Friedman et al., 2015). Consequently, performing model calibration with clinical data is challenging, and the resulting model uncertainty can limit the model’s applicability, e.g., the prediction of an individual dose-response or disease progression (Briggs et al., 2012).

With this work, we aim to address the problem of small and sparse individual time-series data that often poses a bottleneck for calibrating dynamic models with clinical applications. We suggest a Bayesian approach incorporating data sets with observations from many individuals at one sampling time point (cross-sectional data) into the model calibration. The workflow we propose is similar to the method explained in (Tariq et al., 2016). The authors in (Tariq et al., 2016) aim to predict the individual patient outcome by (1.) calibrating a population-average model using the pooled longitudinal data of patients and (2.) “personalizing” the population-average model for a given individual using a Bayesian parameter estimation method. In contrast to (Tariq et al., 2016), we use a cross-sectional population sample to calibrate the population-average model. The advantage of using a cross-sectional population sample is that the data collection is less challenging and time-consuming than time-series data. However, cross-sectional population data does not contain information about individual dynamics. The question we want to address with this work is whether a cross-sectional population data set can be used to construct individual dynamic models.

We demonstrate our approach using a mechanistic model describing the time evolution of three reproductive hormones over the time course of puberty in girls. The model describes a simplified feedback network between these three hormones. The population-average model is calibrated using cross-sectional data of 601 healthy Norwegian girls from the Bergen Growth Study (BGS, 2006; Madsen et al., 2020, 2022). Subsequently, the model analysis guides a reduction of the parameter space and the re-estimation of model parameters in a Bayesian setting (Bayesian updating) to personalize the population-average model.

## 2 Materials and methods

In the following sections, we introduce the mechanistic model used to demonstrate our method, the calibration and analysis steps of the population-average model, and the translation of the population-average model into individual-specific models.

### Mathematical model

During childhood, reproductive hormones such as follicle-stimulating hormone (FSH), luteinizing hormone (LH), and estradiol (E2) are not secreted at concentrations that trigger reproductive function. This lack of reproductive function during childhood is primarily caused by the suppression of the gonadotropin-releasing hormone (GnRH) pulse generator – a functional unit in the brain that regulates the synthesis and release of GnRH (Krsmanovic et al., 2009; Herbison, 2018). The reactivation of the GnRH pulse generator during puberty is key for developing the reproductive system and activating the hypothalamic–pituitary–gonadal axis (HPG axis) (Naulé et al., 2021; Uenoyama et al., 2019; Terasawa, 2022).

The model we introduce describes regulatory feedback loops between GnRH, FSH, LH and E2 (Fig. 1). All are critical elements of the HPG axis and the reproductive system. We encode the reactivation process of the GnRH pulse generator in a semi-mechanistic way by using a sigmoidal input curve for the increase of GnRH release over time,

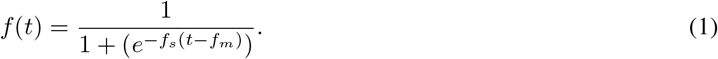

**Figure 1:**
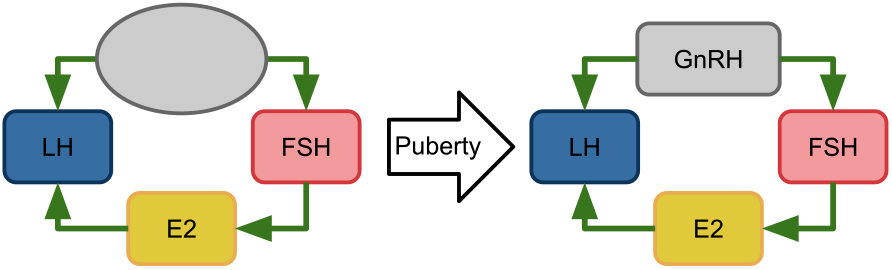
Schematic representation of hormone interactions and re-activation of the HPG axis. With the onset of puberty, the release of GnRH gets enabled. As a result, the signalling along the HPG axis starts working. GnRH stimulates the release of FSH and LH. FSH has a positive feedback effect on E2. E2 again stimulates the release of LH. Additional interactions between those hormones are not represented, because the model neglects them.

The parameter *f*_*s*_ determines the steepness of the curve, i.e., how rapidly the activity increases and the parameter *f*_*m*_ determines the time point at which *f* (*t*) reaches half of its maximum, i.e., the time at which the pulse generator reaches half of its total activity. This formulation is motivated by the observation that GnRH increases gradually and independently of gonadal activity (Terasawa, 2022).

Experimental studies have shown that ovaries can be stimulated before puberty (Wildt et al., 1980; Plant, 2015) and that the HPG axis begins to function when GnRH levels become high enough (Ellison et al., 2012). Those observations motivated us to base the mathematical model formulation of the hormone interactions during pubertal development on a previously developed model for the human menstrual cycle (Fischer-Holzhausen and Röblitz, 2022).

Compared to the menstrual cycle model, the model of the hormone dynamics during female puberty can be drastically reduced in the number and complexity of the ordinary differential equations and parameters. This significant model reduction is possible because the hormone concentrations during puberty do not reach the same peak concentrations as in menstrual cycles. Consequently, we removed all processes from the original model that remained inactive during puberty due to high thresholds of the regulating hormones. Note that the reduced model can, therefore, not be used to simulate late puberty or the first menstrual bleeding (menarche).

A synthesis-clearance relationship describes the dynamics of each hormone. The regulatory interactions (illustrated in Fig. 1) are approximated using Hill functions – a common way to encode feedback interactions as threshold-dependent processes without mechanistic details. Positive Hill functions describe positive feedback actions as sigmoidal curves and are denoted as *H*^+^(*S, T, n*) = *S*^*n*^*/*(*T*^*n*^ + *S*^*n*^). The regulator species *S* regulates another species in a threshold-dependant manner with threshold *T >* 0. The Hill exponent *n >* 0 influences the sigmoidal curve’s steepness and thereby the regulatory process’s rapidity (Santillán, 2008). The proposed model reads as follows:

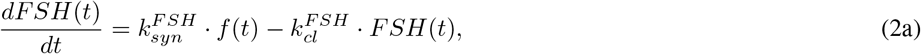

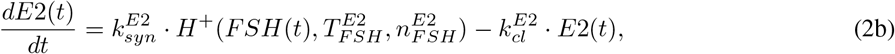

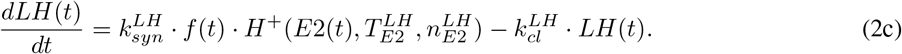

Modulation of the synthesis rate constants 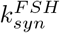 and 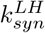 by the function *f* (*t*) in Eqs. (2a) and (2c), respectively, represents the stimulatory action of GnRH on the release of the pituitary hormones FSH and LH (Marques et al., 2022). FSH stimulates the growth of ovarian follicles, hence the release of E2 (Brown, 1978) included in Eq. (2b) via a positive Hill function with FSH concentration threshold 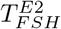 and Hill exponent 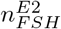. The ovarian hormone E2 has a positive effect on LH (Reed and Carr, 2015), modelled using a positive Hill function with threshold 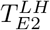 and Hill exponent 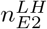

The model is implemented in Python3 (Van Rossum and Drake Jr, 1995), and numerical simulations were performed using SciPy (Virtanen et al., 2020). A model and workflow demonstration is available at https://github.com/SoFiwork/CrossSectional2Individual.

### Experimental data and pre-processing

We use the cross-sectional data set from the Bergen Growth Study 2 (BGS2) (BGS, 2006) to calibrate a population-average model. The BGS2 study includes hormone blood concentrations with chronological age for E2 (No. = 547), LH (No. = 600), and FSH (No. = 599). (Madsen et al., 2020) and (Bruserud et al., 2020) describe the data collection and the cohort composition in detail.

To make the data usable for the calibration of the population-average model, we estimate the continuous mean and standard deviation using the rolling window calculation function provided by the pandas package (Wes McKinney, 2010; The pandas development team, 2020) with a window size of 50 observations.

For the model adaptation to individual time-series data, we use simulated data, because clinical time-series data was unavailable for this work. We simulate time-series data by performing a forward simulation using a parameter set from the stationary parameter distribution of the population-average model, which we obtain from an uncertainty analysis. We add Gaussian noise to the simulated data points and time points to account for noise.

### Population-average model calibration

Figure 2 illustrates our workflow to calibrate and analyse the population-average model. Our routine consists of three main steps:

**Figure 2:**
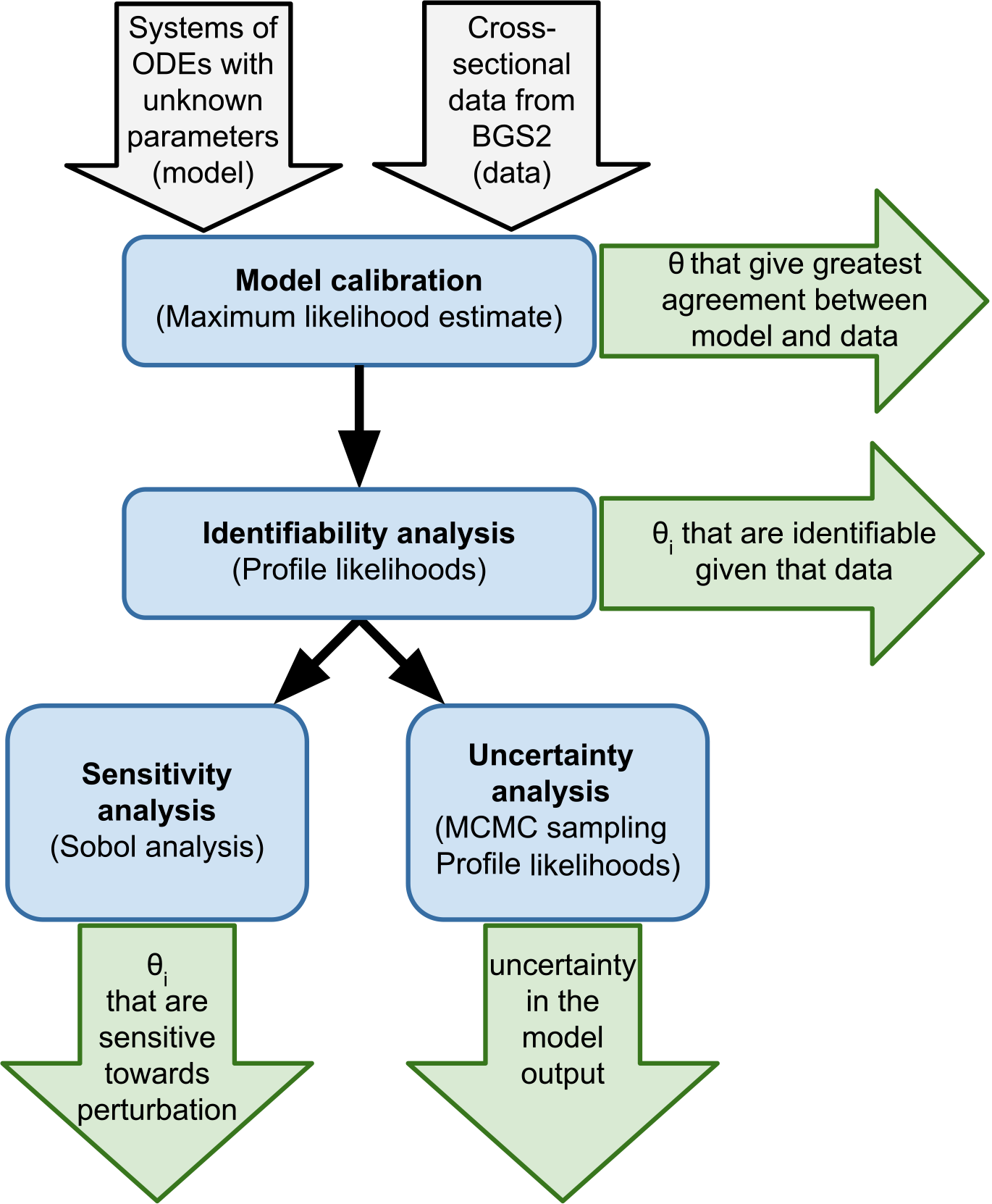
Workflow for calibrating and analysing the population-average model. The pipeline inputs are the mechanistic model and the cross-sectional data set (grey arrows). In the first step, a set of model parameters *θ* that minimizes the distance between the model and the moving average of the cross-sectional data is estimated using ML estimation. In a second step, the likelihood is profiled for each parameter *θ*_*i*_ to identify parameters that can/can not be estimated within bounded confidence intervals. In the third step, the sensitivity and uncertainty of the model are quantified. Sensitivity analysis is performed using the Sobol method. A Markov chain Monte Carlo simulation reveals information about the model uncertainty.

1. Model calibration using maximum likelihood (ML) estimation
2. Identifiability analysis using profile likelihoods (PL)
3. Uncertainty quantification using the Metropolis-Hastings (MH) sampling algorithm and Sobol sensitivity analysis

In the first step, we estimate model parameters *θ* by fitting the model to the processed population-average data *y* using maximum likelihood estimation. The aim is to find a set of parameters 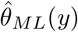 that minimizes the negative log-likelihood. When assuming additive Gaussian measurement noise, this optimisation problem corresponds to minimizing the least squares error between simulated trajectories *x*(*θ*) and associated measurement data *y*, weighted by the estimated standard deviation (Cox and Hinkley, 1979; Eliason, 1993). We use the open-source Python Parameter EStimation TOolbox (pyPESTO) (Schälte et al., 2022) to solve the ML estimation with a multi-start optimization (500 runs). We apply a quasi-Newton method (L-BFGS-B) provided by SciPy (Virtanen et al., 2020) as an optimization algorithm and calculate the gradients numerically with a 3-point finite difference schema provided by the SciPy optimization library (Virtanen et al., 2020). Tab. 1 contains the parameter search regions.

**Table 1:**
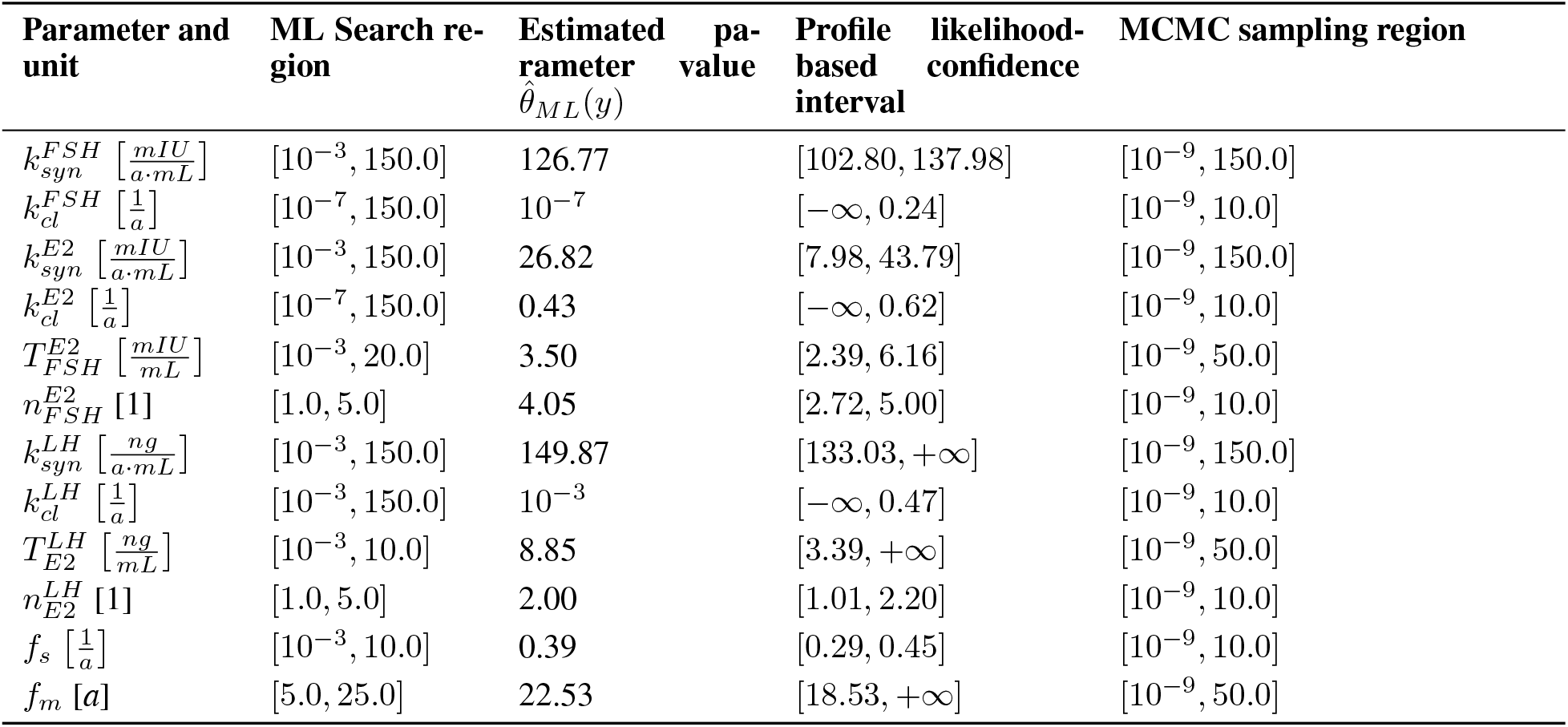
Summary of population-average model parameters. Model parameters are estimated using ML estimation with a search space restricted by the parameter intervals given in the column “Search region”. Confidence intervals result from the analysis of the profiled likelihoods for each parameter.

In the second step, an identifiability analysis is performed by calculating the profile likelihoods (Murphy and Van der Vaart, 2000; Raue et al., 2009) for each model parameter numerically. In doing so, we gain information about ambiguities in the model structure and whether the available data is sufficient to estimate parameter values with finite confidence intervals. In this context, parameters that can not be determined with finite confidence intervals are called non-identifiable. There exist two types of non-identifiable parameters. Structurally non-identifiable parameters are the consequence of redundancies in the model formulation. Practically non-identifiable parameters lack precision because the data do not provide enough information (Wieland et al., 2021; Raue et al., 2009; Kreutz et al., 2013). With a step-wise optimization routine, we obtain a PL for each parameter. Hereby, the parameter *θ*_*i*_ is varied around its optimal value, and for each value of *θ*_*i*_, all remaining parameters *θ*_*j*≠*i*_ are re-optimized using, for example, the ML estimation method. In this work, we use the routine implemented in the pyPESTO package (Schälte et al., 2022) to calculate the profile likelihoods around the optimized model parameters from step one.

In the third step, the model uncertainty is quantified using the Metropolis-Hastings algorithm (Metropolis et al., 1953; Hastings, 1970), to obtain random samples of model parameters according to the likelihood function constructed in step one (Valderrama-Bahamóndez and Fröhlich, 2019). We use an adaptive parallel tempering sampler available in pyPESTO (Schälte et al., 2022) for the MH sampling. Geweke test (Geweke, 1992) is applied to the sampling chain to determine the burn-in period, i.e., the early proportion of the sampling chain that may not converge to the target distribution.

Sensitivity analysis allows studying model output uncertainty resulting from perturbations in the model input (Saltelli et al., 2004). Here we use the Sobol method (Sobol, 1993), which is a variance-based global sensitivity analysis technique. This method quantifies the contribution to the uncertainty in the model output by each parameter individually and by parameter interactions. We use the implementation of Sobol’s sensitivity analysis provided by the SALib package (Herman and Usher, 2017). We decided to allow for parameter perturbations of up to 20% around the estimated parameter value 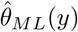.

The results of the model calibration and analysis steps are:

- a parameter point estimate
- identification of identifiable parameters
- identification of sensitive parameters
- distribution and confidence interval of each parameter

We use this information to enable the translation from a population-average model to an individual-specific model, as outlined in the following.

### Individual-specific model calibration

We use a Bayesian parameter estimation framework (Wakefield, 1996; Tariq et al., 2016) to transform the population-average model into an individual-specific model that can be used to predict the individual time-evolution of hormones (Fig. 3). In a Bayesian framework, the posterior distribution *p*(*θ* | *y*) is proportional to the product of the likelihood function *L*(*θ* | *y*), which describes the probability of observing the data *y* given a parameter set *θ*, and the prior distribution *p*(*θ*). Here we use this idea to update our population parameter set (prior parameter distributions *p*(*θ*)) using individual time-series data *y* in the likelihood function to obtain an individual parameter set *p*(*θ* | *y*) as posterior parameter distribution.

**Figure 3:**
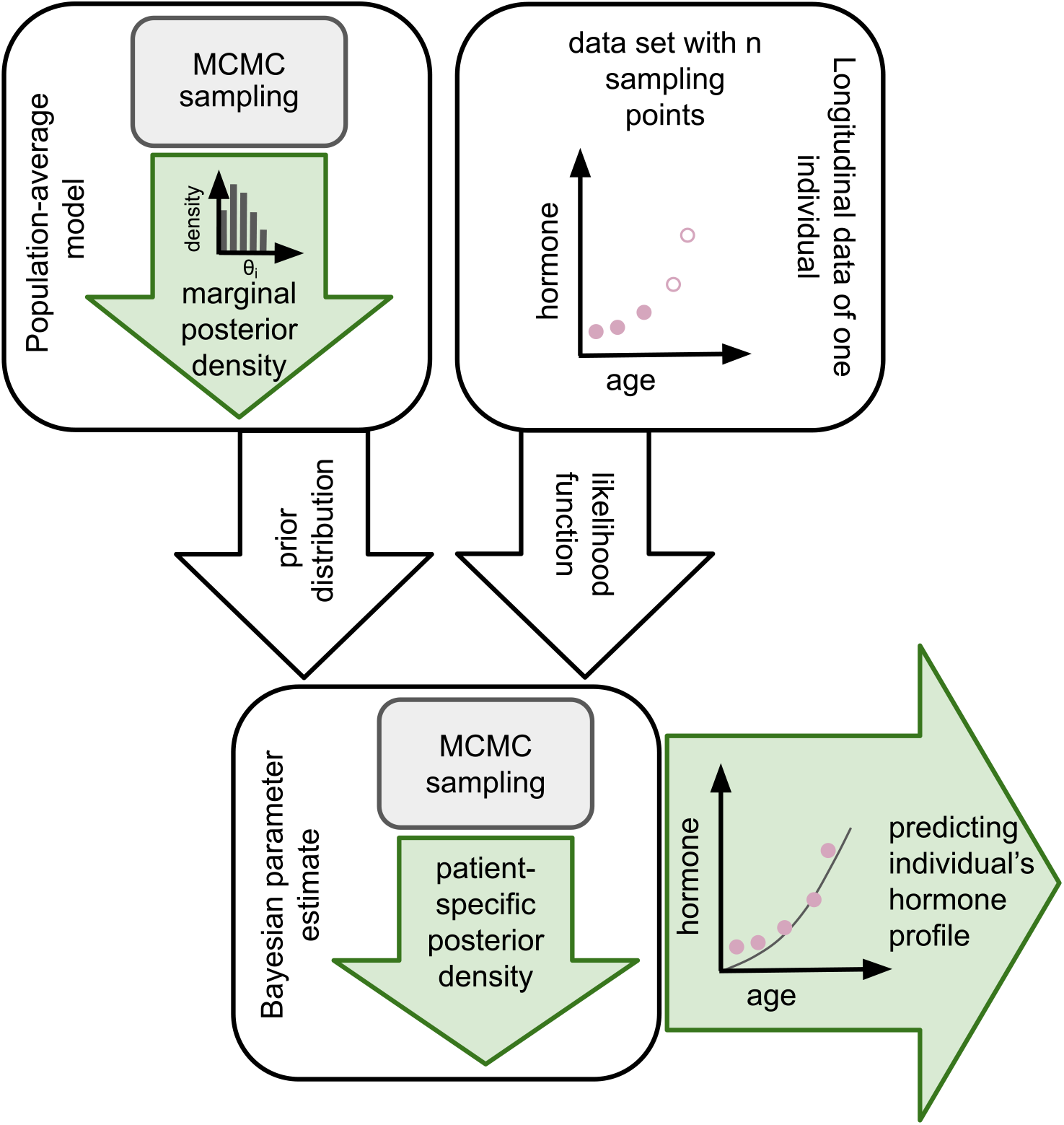
Workflow for Bayesian updating to generate an individual-specific model fit. Individual-specific parameter sets are obtained from a Bayesian parameter estimation approach. The parameter distribution generated with MCMC sampling applied to the population-average model with the cross-sectional data is employed as prior knowledge. For calculating an individual-specific likelihood, only the first 75% of the data points of an individual’s longitudinal data set are used. An individual-specific posterior distribution of the parameters is approximated with MCMC sampling. This parameter distribution can be used for predicting the individual’s hormone dynamics.

In our approach, the prior parameter distribution *p*(*θ*) originates from the uncertainty quantification of the population-average model, where we sampled from the likelihood function using MCMC sampling. Hereby, we obtain histograms of all marginal parameter distributions, to which we then fit continuous log-normal distributions. For the Bayesian parameter estimation, we use the product of these one-dimensional continuous distributions as prior distribution *p*(*θ*). In doing so, we lose information about the correlation between the parameters, but we gain an analytical representation of the priors, from which it is easy to sample.

The likelihood *L*(*θ* | *y*) is based on calculating the mean squared error between the individual time-series data *y* and the simulated model trajectories for a given parameter set *θ*. We solve the Bayesian parameter estimation problem by sampling from the posterior *p*(*θ* | *y*) using the MH algorithm provided by pyPESTO package(Schälte et al., 2022). Hereby, we obtain individual-specific parameter sets. To reduce the complexity of the parameter estimation problem, we do not perform this approach in the full 12-dimensional parameter space but only on those parameters that were identified as sensitive in the Sobol analysis.

## Results and Discussion

We now present the results for calibrating the population-average model, following the workflow in Fig. 2, and its translation into an individual-specific model according to the Bayesian adaptation workflow described in Fig. 3. The data from the Bergen Growth Study 2 is used to calibrate the population-average model, whereas the individual-specific model is calibrated with simulated data. We present the population-average model’s results, followed by the individual-specific model.

### Population-average model

The 12 model parameters were first estimated using ML estimation to find the parameter set that minimizes the error between simulated hormone time-evolution and the averaged hormone concentrations from the BGS2 data. From the multi-start optimization, we selected the parameter set that resulted in the smallest least-squares error between simulated trajectories and data, see Tab. 1. Fig. 4 shows the simulated trajectory with this optimal parameter set. However, the plot of the optimisation history (Fig. A.1 in the Appendix) indicates the existence of several alternative parameter sets which fit the data comparably well.

**Figure 4:**
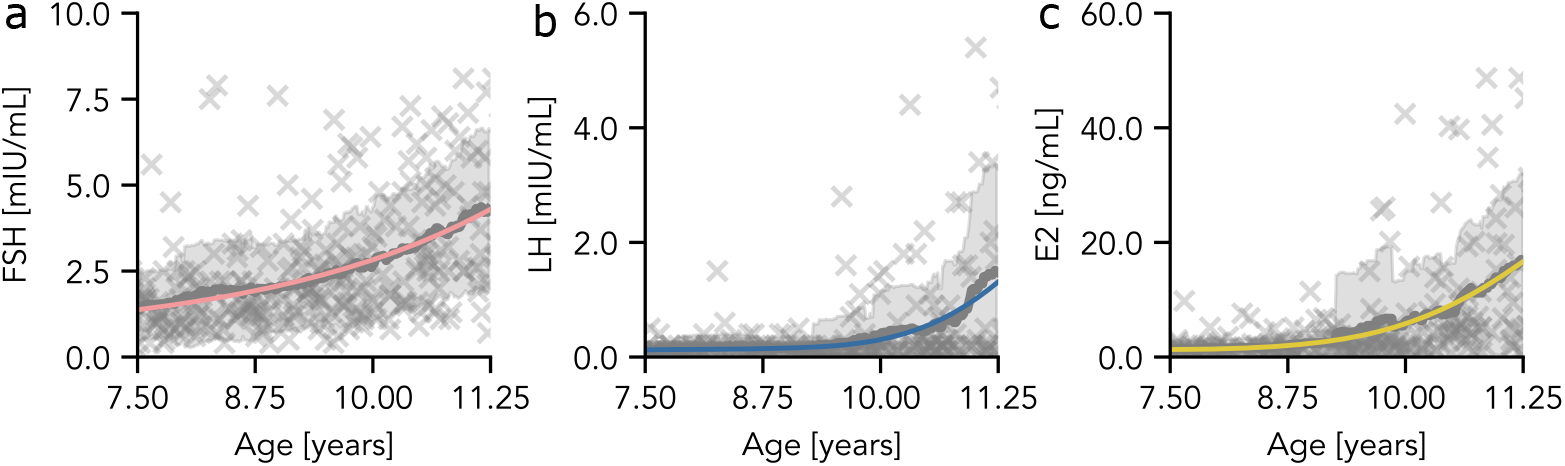
Population-average model simulation results. Representation of the model trajectories of the three sex hormones (FSH, LH, E2) resulting from numerical simulations with the parameter set given in Tab. 1. The moving average is marked as grey lines, and grey areas are the moving standard deviation of the data. The individual measurement points are marked as grey crosses.

Figure 5 represents the PL for each model parameter. The shape of the profile discloses the type of identifiability. Confidence intervals for each parameter (Tab. 1) were derived from the intersection between the profile and the line indicating 95 % confidence interval (Fig. 5). Because all profiles are bounded either towards plus or minus infinity, we can conclude that the model does not have structurally non-identifiable parameters. The parameter profiles presented in Fig. 5(a, c, e, f, j, k) indicate identifiablilty showing finite confidence intervals (Tab. 1). The remaining six parameters are practically non-identifiable. Their confidence intervals miss either lower or upper bounds.

**Figure 5:**
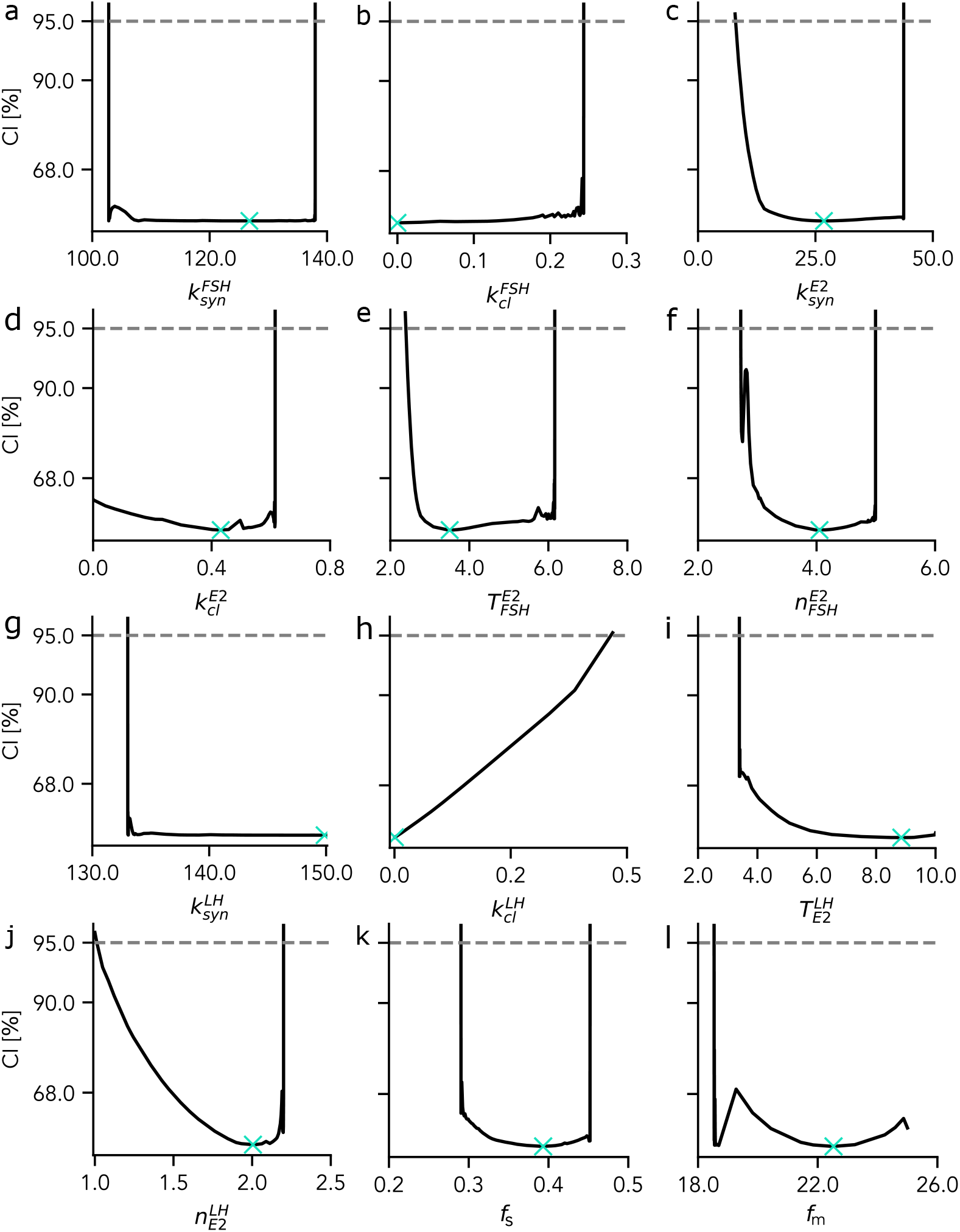
Profile likelihood for each model parameter *θ*_*i*_. The likelihood profile and the ML estimate (marked as a cross) are shown for each parameter separately. The analysis reveals that 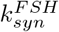 (panel a), 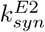 (panel c), 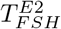 (panel e), 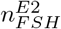 (panel e) 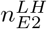 (panel j), and *f*_*s*_ (panel k) are identifiable. In contrast, parameters 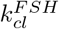 (panel b), 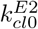 (panel d), 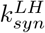 (panel g), 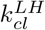 (panel h) 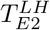 (panel j), and *f*_*m*_ (panel l) are practically non-identifiable.

One way to handle practical non-identifiable parameters is model reduction. The authors in (Tönsing et al., 2018) suggest that parameters with likelihood profiles that flatten out towards minus infinity can be set to zero. We deliberately decided not to apply this approach to our model, because we would remove all clearance terms. The presence of the clearance terms in our model represents the prior biological knowledge, that hormones have a finite lifetime. However, that is not visible in the cross-sectional data. Consequently, we have a model that is not fully identifiable, increasing the uncertainty in the model output.

We performed uncertainty quantification in terms of a full MCMC sampling of the likelihood function (see Tab. 1 for search regions). The marginals of the resulting parameter distribution are illustrated as violin plots in Fig. 6. The ML estimates (Tab. 1) lie within the sampled range (marked as green x) but do not agree with the median or mean of the sampled parameter densities. This is not unexpected because the model is not fully identifiable, and there exist multiple local optima (Raue et al., 2013).

**Figure 6:**
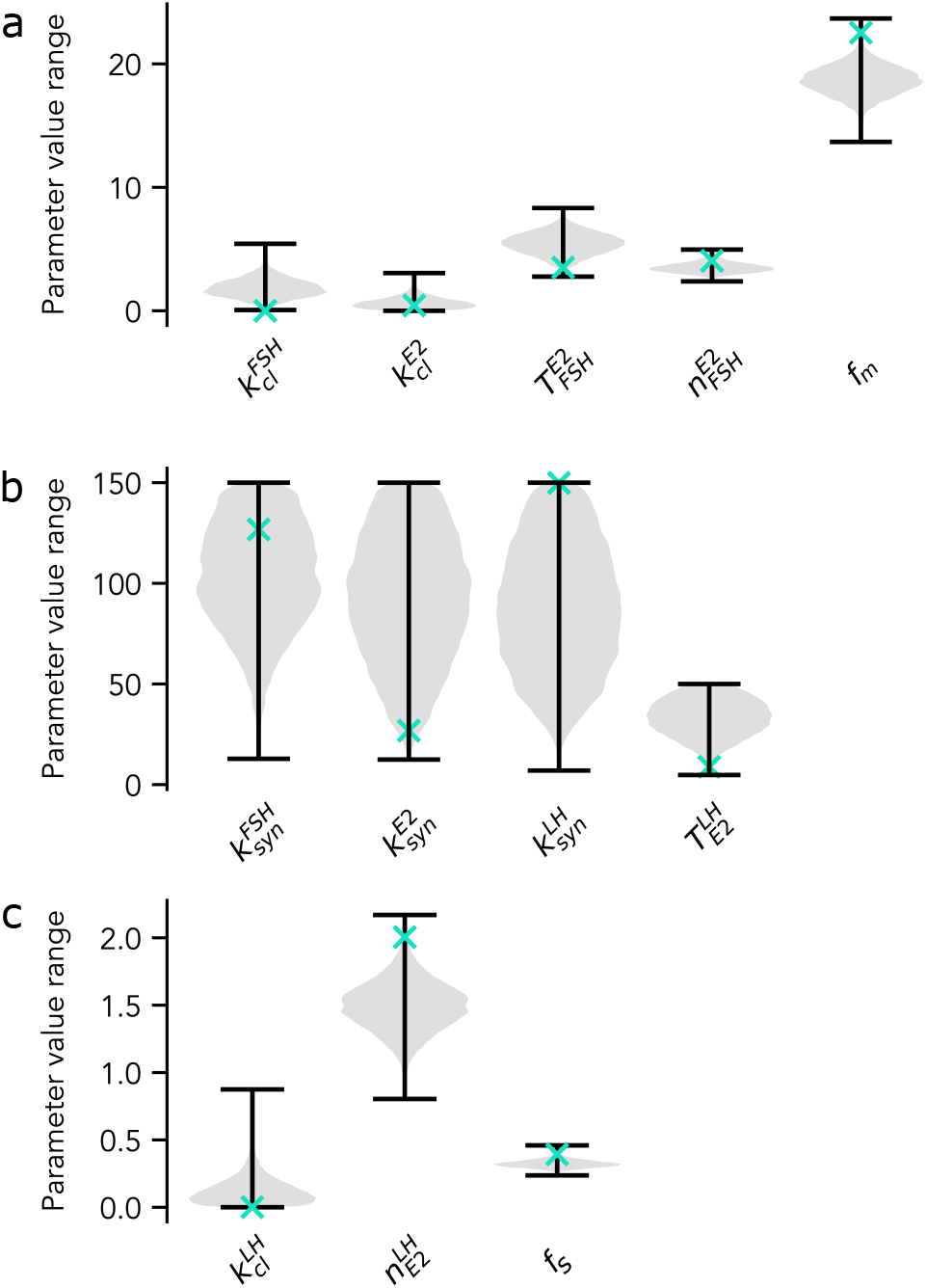
Representation of stationary parameter distributions from MCMC sampling as violin plots. Each violin (grey area) represents a kernel density estimate for each marginal parameter distribution derived from its histogram. Horizontal black lines mark the minimum and maximum of the distributions. Green crosses mark ML estimates.

The Sobol analysis (Fig. 7) reveals that the parameters of the sigmoidal input curve are the most sensitive ones. Additionally, their effect on the uncertainty in the simulation outcome increases with time. Initial values are sensitive during earlier time steps and lose their effect over the simulation time. Other sensitive parameters are 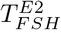 and 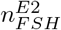, which are both among the identifiable parameters. Perturbations in all other model parameters did not show notable effects on the simulation outcome. Note that sensitivity and practical identifiability are two different concepts because sensitivity does not consider any experimental data. Hence sensitive parameters can be practically non-identifiable (e.g. *f*_*m*_), and practically identifiable parameters do not necessarily need to be sensitive (e.g. 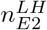).

**Figure 7:**
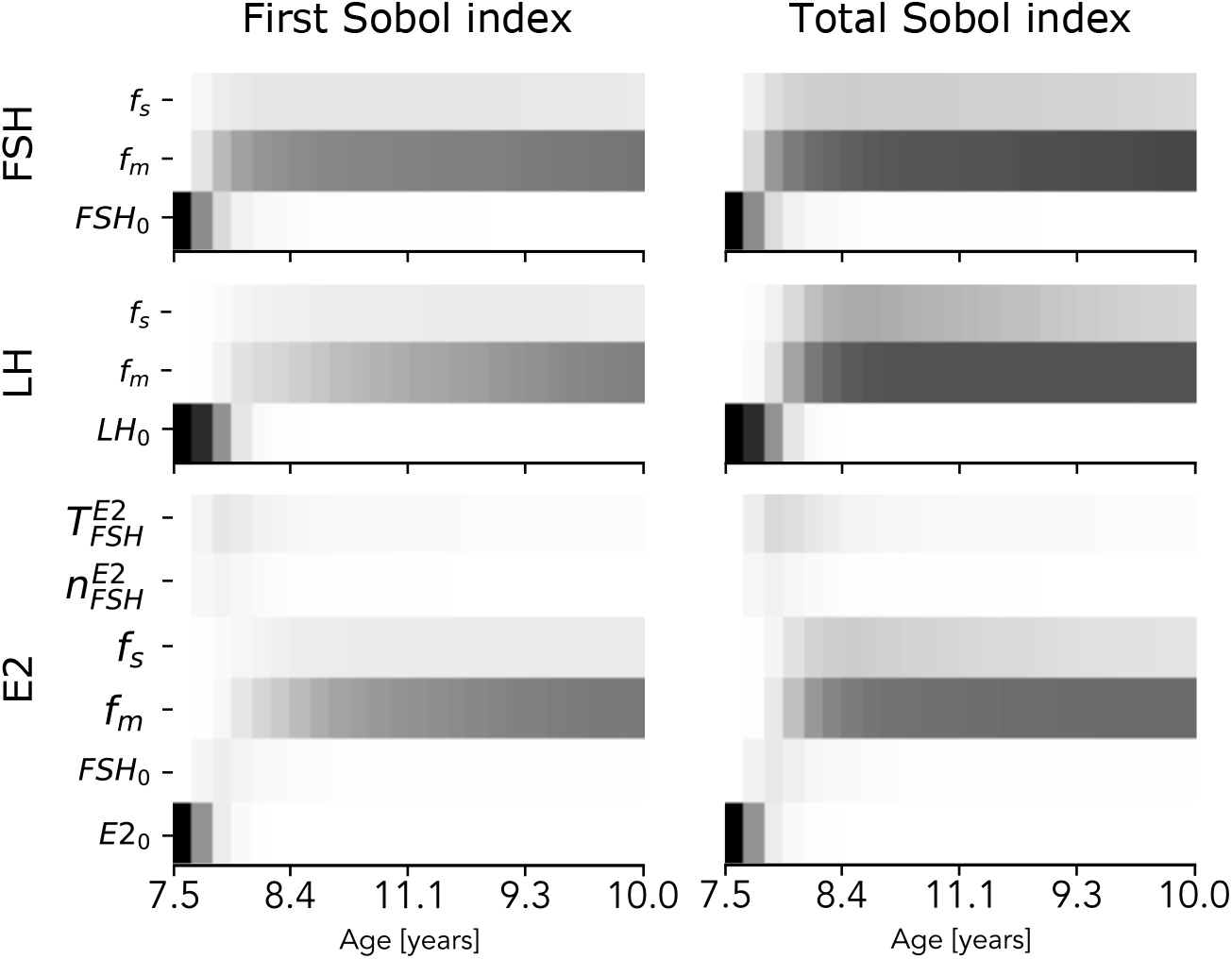
Representation of parameters’ first and total Sobol index over the simulation time. Sensitivity values are encoded as grey scale values (black = highly sensitive and white = not sensitive). For clarity, this figure only represents parameters that are sensitive.

### Individual-specific model

Translating the population-average model to an individual-specific model requires re-estimating model parameters. Based on the Sobol sensitivity analysis (Fig. 7), we selected *f*_*s*_, *f*_*m*_, 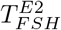, and 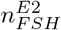 for re-estimation. All other eight parameters remained at the values given in Tab. 1. Reducing the parameter space to sensitive parameters avoids re-estimating parameters that do not affect the model output and thereby increases the efficiency of the MCMC sampling. The initial values of the three hormones were inferred directly from the data as the first data point of the time-series.

For the Bayesian parameter estimation of individual parameter distributions, we not only utilize the results from the sensitivity analysis but also the results from the uncertainty analysis of the population-average model. The insights about the model parameters are used to construct prior parameter distributions. Those prior distributions are updated, resulting in individual posterior distributions with a likelihood solely based on the individual data. By construction, parameter correlations (Fig. A.3 in the Appendix) are neglected by the prior but rediscovered after updating the parameter distributions (Fig. 8).

**Figure 8:**
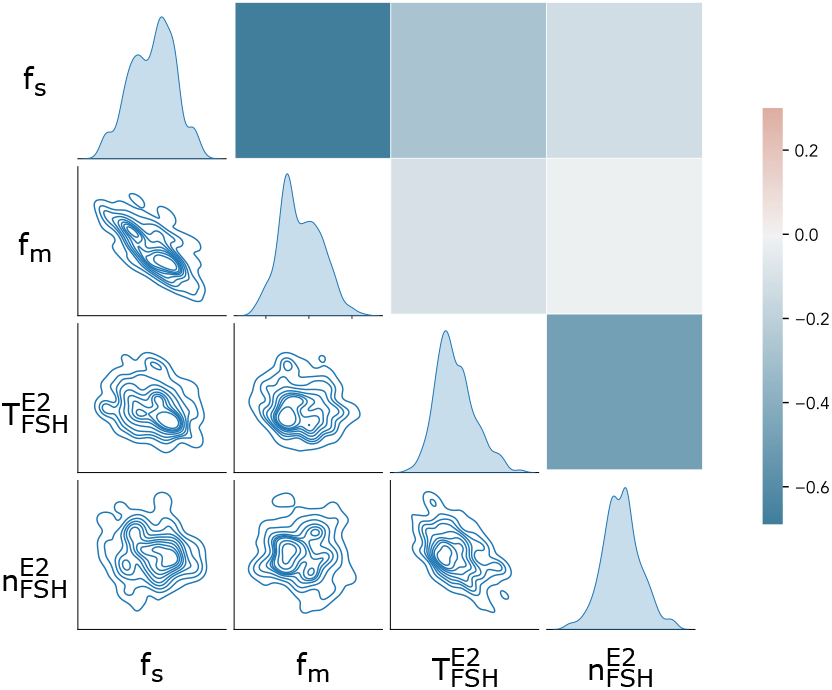
Parameter correlation in the individual-specific model. The upper right half of the matrix contains the pairwise Pearson correlation coefficients. The lower left half shows the corresponding bi-variate densities resulting from marginalization. The marginal parameter distributions are given diagonal.

We used a set of simulated data, generated by a forward simulation with a parameter set from the stationary distribution of the MCMC sampling performed for the population-average model, as an individual data set to demonstrate our adaptation approach. Only one simulated individual is displayed because the range of variability covered by the stationary distribution is not large (see Fig. A.2). The simulation results of the individual-specific model are summarized in Fig. 9. Figs. 9 (a-c) allow for a comparison between the cross-sectional data and the simulated time-series. As expected, the individual data set generated with parameters from the stationary distribution of the population-average model lies within the cross-sectional data set. Figs. 9 (d-f) show simulated trajectories for the individual-specific model. Note that we only used the first three measurement points of the simulated time-series to estimate the individual parameter distributions. The remaining data points can be used to evaluate the model’s predictive performance. All data points, except the last data point of the LH time-series, are within the 99 % confidence interval, generated based on 100000 solutions with parameter sets from the stationary distribution. Figs. 9 (g-j) demonstrate that the population-parameter distributions have been updated for the individual-specific model. For a quantitative comparison between the distributions, the value of the Jensen-Shannon divergence (JSD) between the prior and each posterior is plotted in Fig. 9 (k). It is visible that the posterior distributions of the two parameters from the input curve, *f*_*m*_ and *f*_*s*_, differ more from the prior than the distributions of the Hill function’s parameters 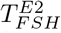 and 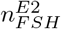. So far, this is only an observation, and further investigations are needed to draw any biological conclusions.

**Figure 9:**
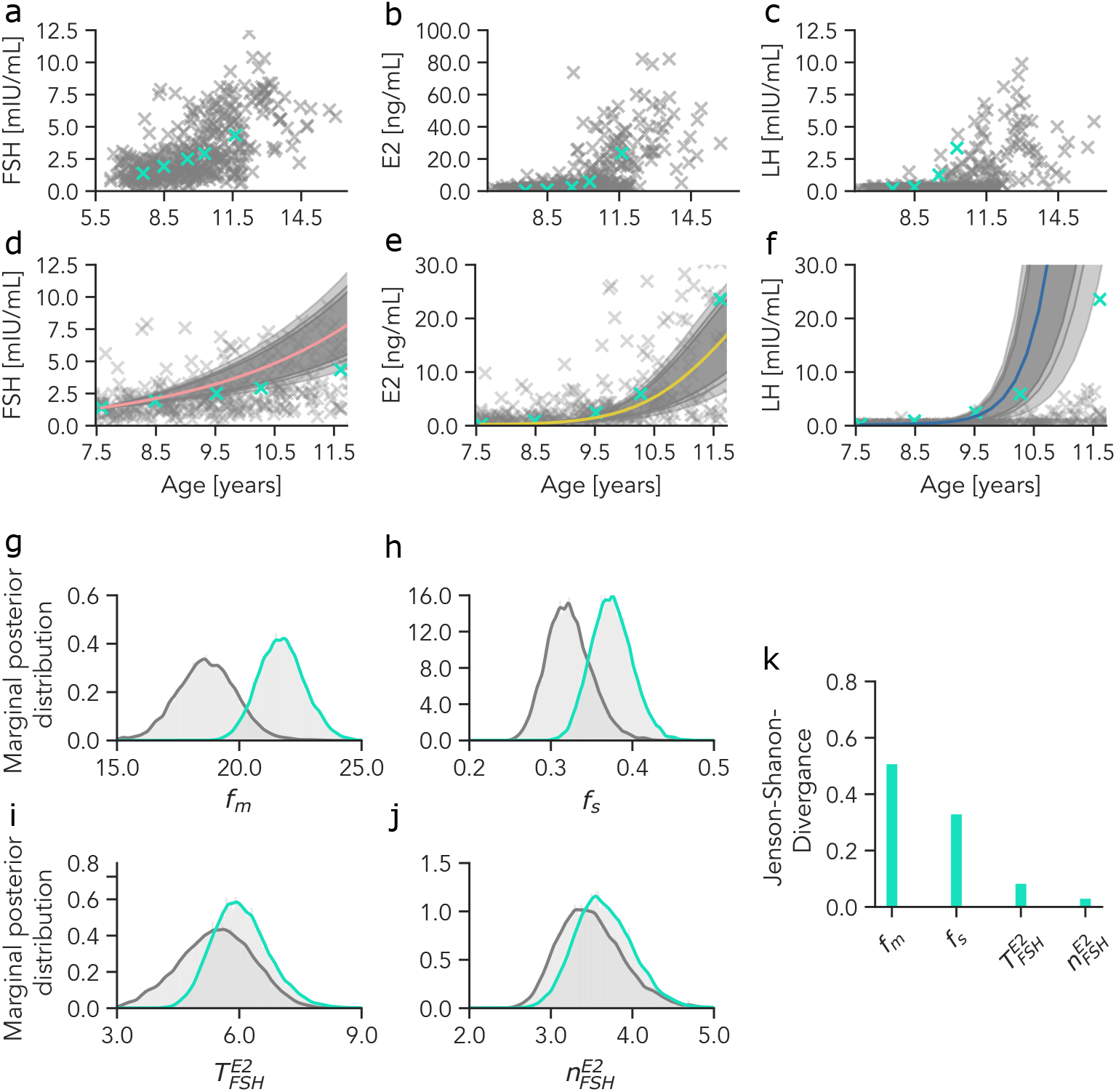
Individual-specific model simulation. Figures (a-c) illustrate the location of simulated time-series data in the cross-sectional data set. Figures (d-f) show the simulated result of the individual-specific model. The presented percentiles (99.0% - light grey, 95.0% - grey, 90.0% - dark grey) are derived from 100000 model simulations with parameter sets from the stationary posterior distribution. Figures (g-j) show the four updated parameters’ marginal distributions (grey - population parameters, green - individual parameters). Those figures indicate that all distributions are updated. Figure k represents the JSD, quantifying the difference between the two distributions for each parameter.

## 3 Conclusion

In this work, we propose a workflow incorporating cross-sectional clinical data into calibration of an individual-specific dynamic model. We demonstrate that a population-average model calibrated with a cross-section data set can be translated into an individual-specific model using Bayesian updating. However, the presented study has some limitations, which we would like to point out in the following.

Using a well-established benchmark model would better underline the value of the proposed workflow. However, we did not have access to a cross-sectional data set for a well-established mechanistic model. We, therefore, decided to construct a mechanistic model for the time evolution of sex hormones during puberty in girls to be able to demonstrate our model calibration pipeline.

The model formulation itself is motivated by biological knowledge (bottom-up approach). Therefore, it would be interesting to compare it to alternative model formulations that fit the data, especially statistical models derived from a top-down approach. One could argue that machine learning (ML) models are expected to have a better predictive performance than mechanistic models. However, ML models are known to be data-hungry and would need training data of high quality and quantity, which are usually not available in a clinical context (Baker et al., 2018). In conclusion, the model formulation was not the central objective of this work, and therefore, we did not investigate alternatives.

From an algorithmic perspective, the construction of the prior distributions for the individual-specific model calibration, as performed in this work, is a subject for discussion. Some kernel density estimation could be applied to the full-dimensional parameter space sample as an alternative to marginalisation. This also results in an analytical representation of the prior from which one can sample, but this representation will depend on the selected bandwidth. Another alternative would be sampling importance re-sampling (SIR), which is based on re-weighting the prior samples. However, this approach assumes that the individual under consideration is sufficiently well-represented by the population-average model given by the prior, which might not always be true. Finally, in a continuous clinical monitoring context, particle filters that allow for sequential data processing are usually more efficient than MCMC methods (Maier et al., 2020).

Another limitation of this work is using simulated longitudinal data instead of clinical data for individual-specific model calibration. Such clinical longitudinal data have been collected, as part of the Copenhagen Puberty Study^1^. In future, we aim to work with these clinical data to systematically assess the predictive performance of individual-specific dynamic models.

Overall, finding individual-specific dynamic models is of interest because they form a basis for further analysis. For example, clustering individual models in the parameter space can help identify phenotypes. We hope that our work advocates the use of Bayesian methods as a means to integrate different data sources into mathematical models.

## Acknowledgment

The authors like to thank Pétur Benedikt Júlíusson and Andre Madsen for providing access to the Bergen Growth Study data.

## A Appendix

### A.1 Optimisation history

Fig. A.1 shows the waterfall plot from the multi-start optimisation to demonstrate the convergence of the optimization runs.

### A.2 Simulation of the population-average model

Fig. A.2 displays percentiles resulting from simulations with sampled parameter sets from the stationary solution of the Metropolis-Hastings sampling routine. It gives an impression of the model uncertainty. For the interpretation of Fig. A.2, one has to keep in mind that the moving average of the data is used for model calibration and sampling and not the cross-sectional data set itself (grey marks in Fig. A.2). As a result, it is not surprising that the percentiles do not cover the entire spread of the BGS 2 data set.

### A.3 Parameter correlations

Fig. A.3 shows the pairwise Pearson correlation coefficients between model parameters together with the bivariate distributions resulting from the MCMC sampling of the likelihood for the population-avarage model.

**Figure A.1:**
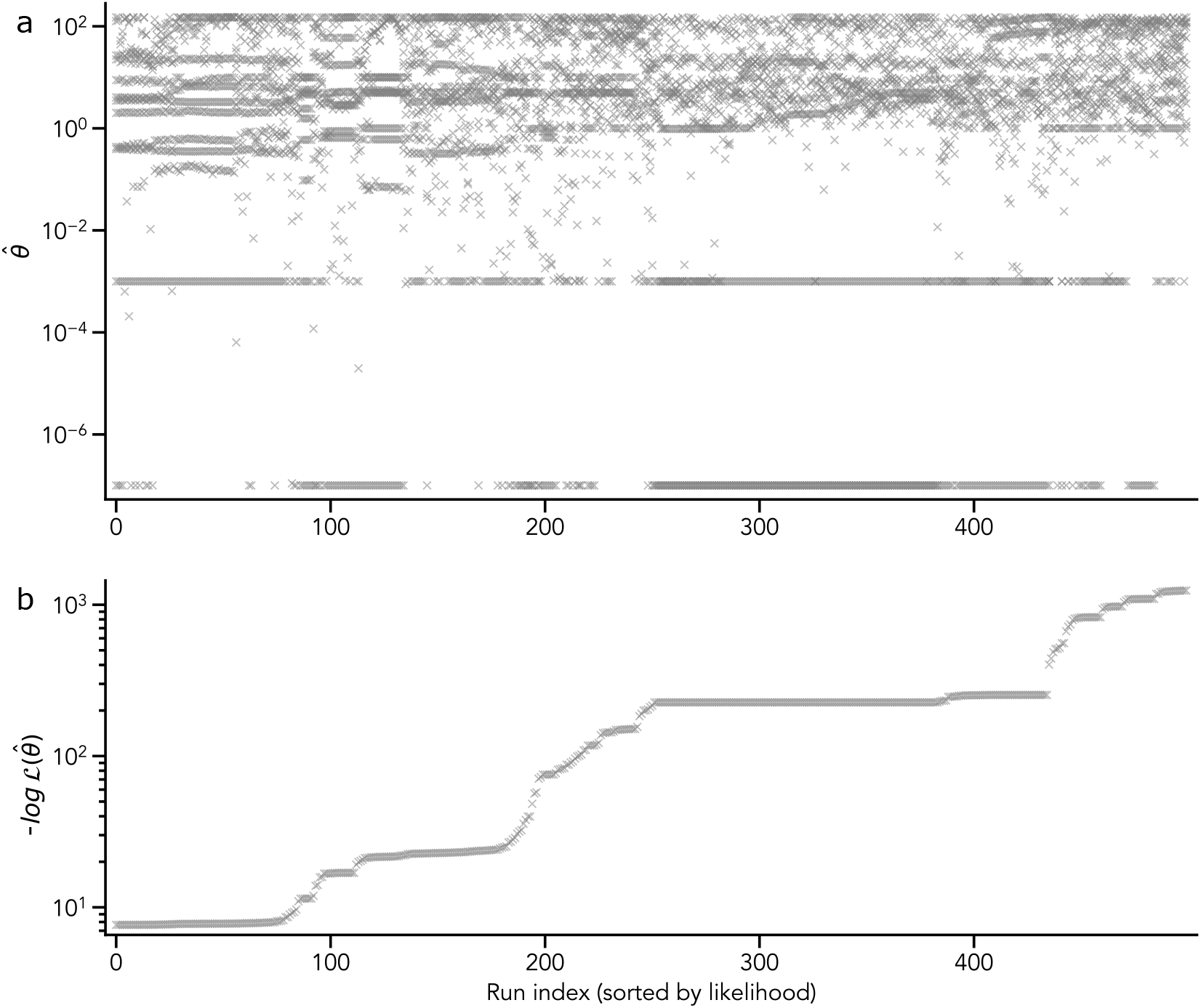
Optimization history. a) Parameter set at the endpoint of each optimisation run. It can be seen that many parameters keep their value in different local optima. b) Values of the negative log-likelihood function at the end of each optimization run.

**Figure A.2:**
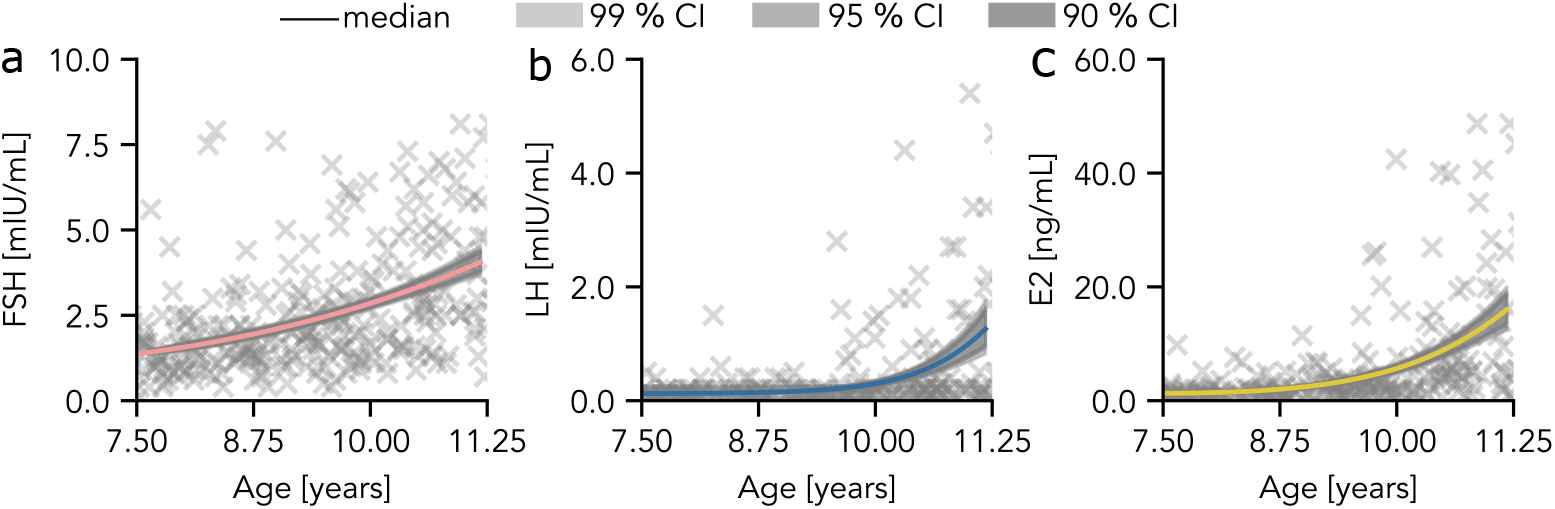
MCMC Sampling based confidence intervals. Illustrated percentiles are derived from model simulations with 150000 parameter sets drawn from the stationary distribution. Confidence intervals (99%, 95%, 90%) are represented as grey areas. The median of the sampling based simulation trajectories is gives as a colored line.

**Figure A.3:**
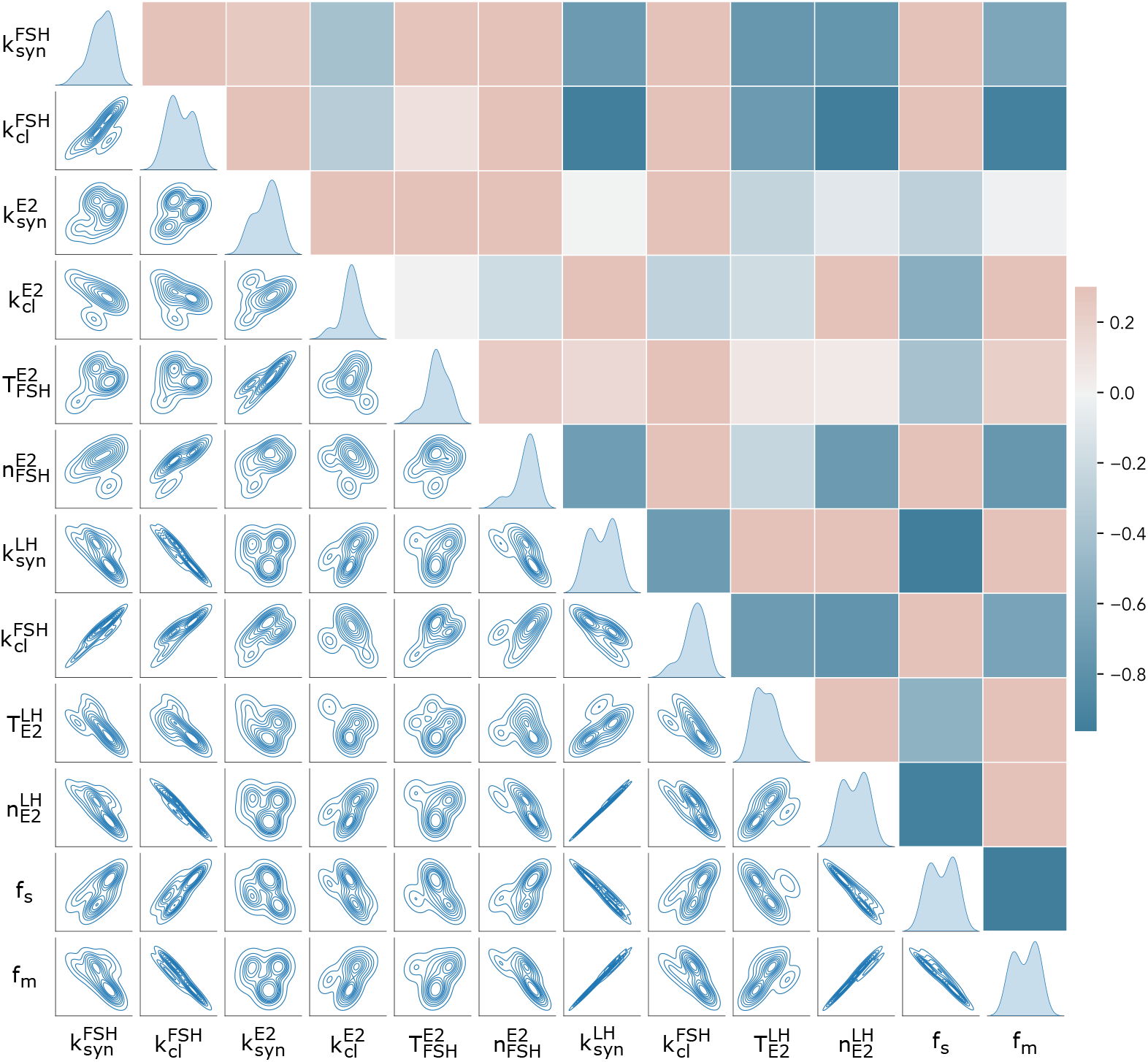
Parameter correlation in the population-average model. The upper right half of the matrix contains the pairwise Pearson correlation coefficients. The lower left half shows the corresponding bi-variate densities resulting from marginalization.

## Footnotes

1 https://www.clinicaltrials.gov/ct2/show/NCT01411527

## References

Baker, R. E., Pena, J.-M., Jayamohan, J., and Jérusalem, A. (2018). Mechanistic models versus machine learning, a fight worth fighting for the biological community? Biology Letters, 14(5):20170660.

BGS (2003-2006). The Bergen Growth Study 1 and 2. https://www.vekststudien.no/en/. Accessed: 06-Oct-2022.

Briggs, A. H., Weinstein, M. C., Fenwick, E. A., Karnon, J., Sculpher, M. J., Paltiel, A. D., Force, I.-S. M. G. R. P. T., et al. (2012). Model parameter estimation and uncertainty: a report of the ISPOR-SMDM modeling good research practices task force-6. Value in Health, 15(6):835–842.

Brown, J. (1978). Pituitary control of ovarian function—concepts derived from gonadotrophin therapy. Australian and New Zealand Journal of Obstetrics and Gynaecology, 18(1):47–54.

Bruserud, I. S., Roelants, M., Oehme, N. H. B., Madsen, A., Eide, G. E., Bjerknes, R., Rosendahl, K., and Juliusson, P. B. (2020). References for ultrasound staging of breast maturation, Tanner breast staging, pubic hair, and menarche in Norwegian girls. The Journal of Clinical Endocrinology & Metabolism, 105(5):1599–1607.

Cox, D. R. and Hinkley, D. V. (1979). Theoretical Statistics. CRC Press.

Eliason, S. R. (1993). Maximum likelihood estimation: Logic and practice, volume 96 of Quantitative Applications in the Social Sciences. Sage.

Ellison, P. T., Reiches, M. W., Shattuck-Faegre, H., Breakey, A., Konecna, M., Urlacher, S., and Wobber, V. (2012). Puberty as a life history transition. Annals of Human Biology, 39(5):352–360.

Fischer-Holzhausen, S. and Röblitz, S. (2022). Hormonal regulation of ovarian follicle growth in humans: Model-based exploration of cycle variability and parameter sensitivities. Journal of Theoretical Biology, page 111150.

Friedman, L. M., Furberg, C. D., DeMets, D. L., Reboussin, D. M., and Granger, C. B. (2015). Fundamentals of Clinical Trials. Springer.

Gábor, A. and Banga, J. R. (2015). Robust and efficient parameter estimation in dynamic models of biological systems. BMC Systems Biology, 9(1):1–25.

Geweke, J. (1992). Evaluating the accuracy of sampling-based approaches to the calculations of posterior moments. Bayesian Statistics, 4:641–649.

Greenland, S., Mansournia, M. A., and Altman, D. G. (2016). Sparse data bias: a problem hiding in plain sight. BMJ, 352.

Hastings, W. K. (1970). Monte Carlo sampling methods using Markov chains and their applications. Biometrika, 57(1):97–109.

Herbison, A. E. (2018). The gonadotropin-releasing hormone pulse generator. Endocrinology, 159(11):3723–3736.

Herman, J. and Usher, W. (2017). SALib: An open-source python library for sensitivity analysis. The Journal of Open Source Software, 2(9).

Kreutz, C., Raue, A., Kaschek, D., and Timmer, J. (2013). Profile likelihood in systems biology. The FEBS journal, 280(11):2564–2571.

Krsmanovic, L. Z., Hu, L., Leung, P.-K., Feng, H., and Catt, K. J. (2009). The hypothalamic GnRH pulse generator: multiple regulatory mechanisms. Trends in Endocrinology & Metabolism, 20(8):402–408.

Madsen, A., Almås, B., Bruserud, I. S., Oehme, N. H. B., Nielsen, C. S., Roelants, M., Hundhausen, T., Ljubicic, M. L., Bjerknes, R., Mellgren, G., et al. (2022). Reference curves for pediatric endocrinology: leveraging biomarker Z-scores for clinical classifications. The Journal of Clinical Endocrinology & Metabolism, 107(7):2004–2015.

Madsen, A., Bruserud, I. S., Bertelsen, B.-E., Roelants, M., Oehme, N. H. B., Viste, K., Bjerknes, R., Almås, B., Rosendahl, K., Mellgren, G., et al. (2020). Hormone references for ultrasound breast staging and endocrine profiling to detect female onset of puberty. The Journal of Clinical Endocrinology & Metabolism, 105(12):e4886–e4895.

Maier, C., Hartung, N., de Wiljes, J., Kloft, C., and Huisinga, W. (2020). Bayesian data assimilation to support informed decision making in individualized chemotherapy. CPT: Pharmacometrics & Systems Pharmacology, 9(3):153–164.

Marques, P., Skorupskaite, K., Rozario, K. S., Anderson, R. A., and George, J. T. (2022). Physiology of GnRH and gonadotropin secretion. Endotext [Internet].

Metropolis, N., Rosenbluth, A. W., Rosenbluth, M. N., Teller, A. H., and Teller, E. (1953). Equation of state calculations by fast computing machines. The Journal of Chemical Physics, 21(6):1087–1092.

Motta, S. and Pappalardo, F. (2013). Mathematical modeling of biological systems. Briefings in Bioinformatics, 14(4):411–422.

Murphy, S. A. and Van der Vaart, A. W. (2000). On profile likelihood. Journal of the American Statistical Association, 95(450):449–465.

Naulé, L., Maione, L., and Kaiser, U. B. (2021). Puberty, a sensitive window of hypothalamic development and plasticity. Endocrinology, 162(1):bqaa209.

Plant, T. M. (2015). 60 years of neuroendocrinology: The hypothalamo-pituitary–gonadal axis. Journal of Endocrinology, 226(2):T41–T54.

Raue, A., Kreutz, C., Maiwald, T., Bachmann, J., Schilling, M., Klingmüller, U., and Timmer, J. (2009). Structural and practical identifiability analysis of partially observed dynamical models by exploiting the profile likelihood. Bioinformatics, 25(15):1923–1929.

Raue, A., Kreutz, C., Theis, F. J., and Timmer, J. (2013). Joining forces of Bayesian and frequentist methodology: a study for inference in the presence of non-identifiability. Philosophical Transactions of the Royal Society A: Mathematical, Physical and Engineering Sciences, 371(1984):20110544.

Reed, B. G. and Carr, B. R. (2015). The normal menstrual cycle and the control of ovulation. Endotext [Internet]. Available from: https://www.ncbi.nlm.nih.gov/books/NBK279054/.

Robinson, K. A., Dennison, C. R., Wayman, D. M., Pronovost, P. J., and Needham, D. M. (2007). Systematic review identifies number of strategies important for retaining study participants. Journal of Clinical Epidemiology, 60(8):757–e1.

Saltelli, A., Tarantola, S., Campolongo, F., Ratto, M., et al. (2004). Sensitivity analysis in practice: a guide to assessing scientific models. Chichester, England.

Santillán, M. (2008). On the use of the Hill functions in mathematical models of gene regulatory networks. Mathematical Modelling of Natural Phenomena, 3(2):85–97.

Schälte, Y., Fröhlich, F., Stapor, P., Vanhoefer, J., Weindl, D., Jost, P. J., Wang, D., Lakrisenko, P., Raimúndez, E., Pathirana, D., Schmiester, L., Städter, P., Contento, L., Merkt, S., Dudkin, E., Grein, S., and Hasenauer, J. (2022). pyPESTO - Parameter EStimation TOolbox for python, version v0.2.14. Zenodo. https://doi.org/10.5281/zenodo.7248648.

Sebastian-Gambaro, M. A., Liron-Hernandez, F. J., and Fuentes-Arderiu, X. (1997). Intra-and inter-individual biological variability data bank. European Journal of Clinical Chemistry and Clinical Biochemistry, 35:845–852.

Sobol, I. M. (1993). Sensitivity analysis for non-linear mathematical models. Mathematical Modelling and Computational Experiment, 1:407–414.

Tariq, I., Chen, T., Kirkby, N. F., and Jena, R. (2016). Modelling and Bayesian adaptive prediction of individual patients’ tumour volume change during radiotherapy. Physics in Medicine & Biology, 61(5):2145.

Terasawa, E. (2022). The mechanism underlying the pubertal increase in pulsatile GnRH release in primates. Journal of Neuroendocrinology, page e13119.

The pandas development team (2020). pandas-dev/pandas: Pandas. Zenodo. https://doi.org/10.5281/zenodo.3509134.

Tönsing, C., Timmer, J., and Kreutz, C. (2018). Profile likelihood-based analyses of infectious disease models. Statistical methods in medical research, 27(7):1979–1998.

Udtha, M., Nomie, K., Yu, E., and Sanner, J. (2015). Novel and emerging strategies for longitudinal data collection. Journal of Nursing Scholarship, 47(2):152–160.

Uenoyama, Y., Inoue, N., Nakamura, S., and Tsukamura, H. (2019). Central mechanism controlling pubertal onset in mammals: A triggering role of kisspeptin. Frontiers in Endocrinology, 10:312.

Valderrama-Bahamóndez, G. I. and Fröhlich, H. (2019). MCMC techniques for parameter estimation of ODE based models in systems biology. Frontiers in Applied Mathematics and Statistics, 5:55.

Van Rossum, G. and Drake Jr, F. L. (1995). Python reference manual. Centrum voor Wiskunde en Informatica Amsterdam.

Villaverde, A. F., Pathirana, D., Fröhlich, F., Hasenauer, J., and Banga, J. R. (2022). A protocol for dynamic model calibration. Briefings in Bioinformatics, 23(1):bbab387.

Virtanen, P., Gommers, R., Oliphant, T. E., Haberland, M., Reddy, T., Cournapeau, D., Burovski, E., Peterson, P., Weckesser, W., Bright, J., van der Walt, S. J., Brett, M., Wilson, J., Millman, K. J., Mayorov, N., Nelson, A. R. J., Jones, E., Kern, R., Larson, E., Carey, C. J., Polat, İ., Feng, Y., Moore, E. W., VanderPlas, J., Laxalde, D., Perktold, J., Cimrman, R., Henriksen, I., Quintero, E. A., Harris, C. R., Archibald, A. M., Ribeiro, A. H., Pedregosa, F., van Mulbregt, P., and SciPy 1.0 Contributors (2020). SciPy 1.0: Fundamental Algorithms for Scientific Computing in Python. Nature Methods, 17:261–272.

Wakefield, J. (1996). Bayesian individualization via sampling-based methods. Journal of Pharmacokinetics and Biopharmaceutics, 24(1):103–131.

Wes McKinney (2010). Data Structures for Statistical Computing in Python. In Stéfan van der Walt and Jarrod Millman, editors, Proceedings of the 9th Python in Science Conference, pages 56–61.

Wieland, F.-G., Hauber, A. L., Rosenblatt, M., Tönsing, C., and Timmer, J. (2021). On structural and practical identifiability. Current Opinion in Systems Biology, 25:60–69.

Wildt, L., Marshall, G., and Knobil, E. (1980). Experimental induction of puberty in the infantile female rhesus monkey. Science, 207(4437):1373–1375.

